# Design of Cyclic Peptides Targeting Protein-Protein Interactions using AlphaFold

**DOI:** 10.1101/2023.08.20.554056

**Authors:** Takatsugu Kosugi, Masahito Ohue

## Abstract

More than 930,000 protein-protein interactions (PPIs) have been identified in recent years, but their physicochemical properties differ from conventional drug targets, complicating the use of conventional small molecules as modalities. Cyclic peptides are a promising modality for targeting protein-protein interactions (PPIs), but it is difficult to predict the structure of a target protein-cyclic peptide complex or to design a cyclic peptide sequence that binds to the target protein using computational methods. Recently, AlphaFold with a cyclic offset has enabled predicting the structure of cyclic peptides, thereby enabling de novo cyclic peptide designs. We developed a cyclic peptide complex offset to enable the structural prediction of target proteins and cyclic peptide complexes and found AlphaFold2 with a cyclic peptide complex offset can predict structures with high accuracy. We also applied the cyclic peptide complex offset to the binder hallucination protocol of AfDesign, a de novo protein design method using AlphaFold, and we could design a high predicted local-distance difference test and lower separated binding energy per unit interface area than the native MDM2/p53 structure. Furthermore, the method was applied to 12 other protein-peptide complexes and one protein-protein complex. Our approach shows that it is possible to design putative cyclic peptide sequences targeting PPI.

## 1. Introduction

Protein-protein interactions (PPIs) play a crucial role in numerous biological and biochemical processes, including signal transduction and metabolism in cellular activities [1]. Over 930,000 human PPIs have been identified to date [2] (BioGrid 4.4.222, June 2023). Since the early 2000s, PPIs have garnered significant attention as potential drug targets for human diseases [3–7]. However, the physicochemical properties of PPI drug targets are distinct from conventional drug targets, presenting challenges in PPI-targeting drug discovery [8,9]. Cyclic peptides have emerged as more effective PPI drug target modalities compared to small-molecule compounds, boasting advantages in stability, structure, and membrane permeability [10–12].

- Due to the absence of termini, cyclic peptides are more resistant to digestive enzymes like peptidases and exoproteases.
- The constraint of the cyclic structure facilitates more stable folding without relying on secondary structures.
- Cyclic structures can also be engineered for permeability across cell membranes, targeting intracellular PPIs that are inaccessible to large molecules or antibodies [13– 16].

There exist several methodologies for the experimental design of PPI-targeting peptides, such as mRNA display and cDNA display, which are used to screen a vast number of0020peptides for protein interactions, however, these methods are prohibitively expensive [17– 19]. Thus, a method for computationally designing candidate peptide sequences that engage with target proteins is much sought after. While there have been instances of designing sequences with a specific function, like antimicrobial peptide sequences [20,21], they require a significant amount of experimental data. Acquiring such extensive data on peptide sequences interacting with specific PPI-targeting proteins is challenging. Computational methods exist to generate peptides similar to known PPI inhibitors [22], but there are few small molecules that can inhibit PPIs [7,23].

High-throughput virtual screening (HTVS) technology is a potent and efficient method for pinpointing drug candidates from a vast compound library [24,25]. Thus, conducting HTVS of peptides with the right virtual screening steps and strategies could feasibly yield bioactive peptide hit candidates [26]. The primary technology underpinning HTVS is molecular docking tools, which encompass both commercial and open-source tools. There are commercial options available, such as the Molecular Operating Environment suite, the Schr”odinger modeling suite, and the OpenEye Scientific Software suite. Tools like Schr”odinger Glide can perform protein-peptide docking, though they’re essentially crafted for linear peptides [27,28]. When these tools are employed for cyclic peptides, the peptide is treated as a ligand, and protein-ligand docking is undertaken. Glide has introduced a macrocycle module that accommodates both flexible and rigid docking [29]. Nevertheless, these tools are optimized primarily for conventional small molecule compounds. A thorough understanding of the parameters for conformation generation and scoring functions is indispensable when docking larger cyclic peptides as ligands.

Conversely, while numerous open-source tools exist for global protein-peptide docking, such as CABS-dock [30,31], ClusPro PeptiDock [32], and PIPER-FlexPepDock [33], and for local protein-peptide docking, including PEP-FOLD3 [34] and DINC 2.0 [35], they cannot handle cyclic peptides. Only two tools, HADDOCK 2.4 [36] and AutoDock CrankPep (ADCP)[37], are specifically tailored for protein-cyclic peptide docking, with ADCP being recognized as the state-of-the-art. Despite significant advancements in peptide docking tools, which are crucial for peptide HTVS strategies, the pace of progress in peptide HTVS remains considerably slower than that for small molecule compounds. Present HTVS methodologies, even those utilizing cutting-edge techniques, appear insufficient in fully tapping the potential of peptide libraries. Notably, few reports in the literature highlight effective use of HTVS, with only a handful of studies showcasing success through both proprietary and open computational techniques [38].

Given this context, we introduce a novel method in this study to design cyclic peptides targeting PPI. This method harnesses a deep learning (DL) based protein structure prediction, diverging from the traditional HTVS strategies reliant on molecular docking. DL techniques, notably AlphaFold2 (AF2) and RoseTTAFold, have made significant strides in the accurate prediction of 3D protein structures from amino acid sequences [39–41]. These tools have demonstrated capabilities in predicting individual proteins as well as more complex assemblies like protein complexes and protein-peptide complexes [42–45]. AF2 can distinguish between favorable and unfavorable conformational templates. This suggests that AF2 learns the approximate biophysical energy function of energetically stable protein backbones, and that DL-based structure prediction methods can link the sequence space and protein structure space to explore protein structures that satisfy a given design [46]. Deep network hallucinations, a DL-based design method, can address a variety of protein designs by starting with random and iterative updating of the protein sequence until the desired properties specified in the loss function are obtained [47]. To increase the possibility of generating more accurate predictions, the loss function converges more quickly using sequence gradients obtained by inverting the structure prediction network and using continuous logits, rather than discrete one-hot encodings of the sequence representation [48–50]. In our previous study, we designed linear peptide binders in AfDesign using a three-step sequence representation of logits, softmax, and one-hot steps [51]. AfDesign binder hallucination is based on AF2, so in principle it is not possible to design cyclic peptides. However, if AF2 can predict he structure of protein-cyclic peptide complexes, it can design sequences that are likely to form complexes with the target protein, which is quite efficient methods.

Recently, using cyclic offset as the relative positional encoding in AF2 has enabled the prediction of head-to-tail cyclic peptides with high accuracy and the design of cyclic peptides “by itself” [52]. However, no DL-based approach has been used to predict protein-cyclic peptide complexes or to design cyclic peptide binders targeting PPIs. Therefore, this study aimed to demonstrate the feasibility of designing a cyclic peptide binder that targets the PPIs of protein-peptide complexes. We first developed relative positional encoding for protein-cyclic peptide complexes, and the predictions were more accurate than those of state-of-the-art local docking tools for cyclic peptide complexes. In this study, the MDM2/p53 complex, a known cancer suppressor gene, was selected as a representative example of a PPI protein-protein complex from the aspect of drug discovery. To evaluate whether it can be applied in various types of protein-peptide complexes, based on previous studies of protein-peptide structure prediction and peptide design, 12 protein-peptide complexes for which AF2 can predict root mean square deviation (RMSD) within 2 Å were selected. For those complexes, the same cyclic offset was applied in AfDesign binder hallucination to design cyclic peptides.

## 2. Materials and Methods

### 2.1. ColabFold settings

Protein-peptide complex structure prediction was performed using the publicly available ColabFold with the following modifications to the localcolabfold installation [53]. An offset matrix for the relative positional encoding of a target protein and cyclic peptide complex prediction was developed by modifying `batch.py` in ColabFold implementation as follows:

1. To predict target protein-peptide complexes, we first used the target protein and cyclic peptide sequences connected by “:” and input to colab_batch as “TARGETPROTEIN-SEQ:CYCLICPEPTIDESEQ”.
2. Let the last residue number of the target protein residue_index + 50 be the first value of the cyclic peptide residue_index (this is known as a chain break, a method for complex prediction [53,54])
3. Using the chain brake residue_index, an offset matrix was created using the default method of AlphaFold-Multimer.
4. A cyclic offset matrix was created for the length of the cyclic peptide in the same way as in the study [52], note that the fixed version of the offset matrix was used here [55].
5. Using the offset matrix for the default complex created in step 3 and the offset matrix for the cyclic peptide created in step 4, we replaced cyclic offsets only in the default offset matrix that corresponds to the cyclic peptide (Figure 1**a**). This enables the prediction of complexes of protein and cyclic peptide, where target proteins are linearized and peptides are cyclized.

**Figure 1.**
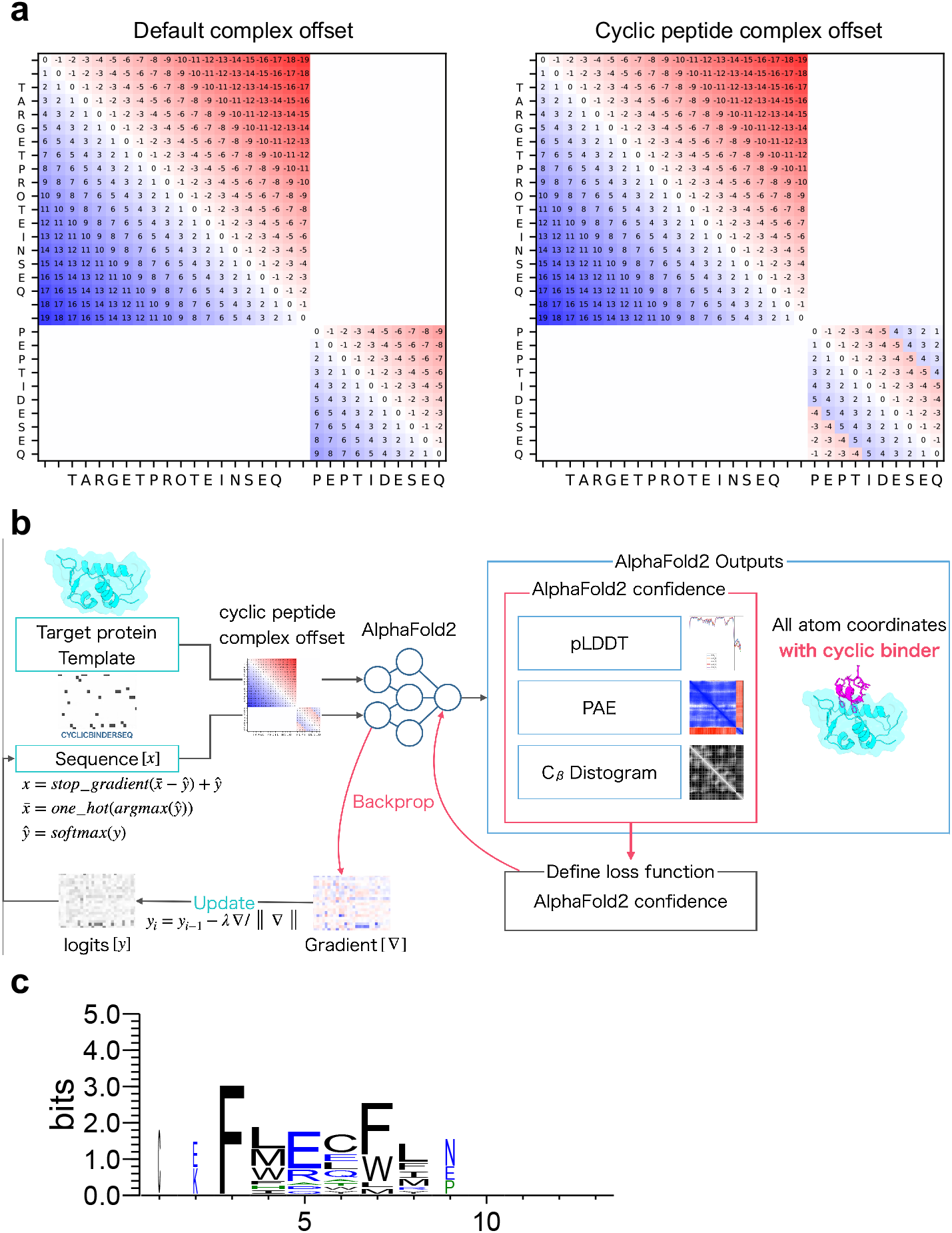
Schematic diagram of cyclic peptide binder hallucination. (**a**) Example relative position encoding of a hypothetical 20-residue protein and a 10-residue peptide. Default complex offset on the left and cyclic peptide complex offset on the right. (**b**) Schematic showing the optimization of the cyclic peptide binder hallucination. The AF2 output is an all-atom coordinate, two reliability indices (PAE and pLDDT), and a distogram. These are used to define the loss function, which is back-propagated to compute the gradient for the design sequence, then updated and predicted in a loop for optimization. A target protein structure and an initial random sequence are input, and a cyclic peptide is hallucinated to the target protein via the cyclic peptide complex offset. (**c**) Example sequence logo for a cyclic peptide design targeting MDM2 considering 3D coordinates. The sequence logo represents each column of alignment as a stack of letters, with the height of each letter proportional to the observed frequency of the corresponding amino acid, and the overall height of each stack proportional to the degree of sequence conservation at that location. The width of the letters is proportional to ungaps of each column. Hydrophilic amino acids (R, K, D, E, N, and Q) are shown in blue, neutral amino acids (S, G, H, T, A, and P) are shown in green, and hydrophobic amino acids (Y, V, M, C, L, F, I, and W) are shown in black.

The sequence search option for building multiple sequence alignment (MSA) was specified as MMseqs2 (mmseqs_uniref_env). When predicting protein-cyclic peptide complex structure registered in the Protein Data Bank (PDB), we used alphafold2_multimer_v2 model (AF_v2), considering the length cutoff of the AF2 training dataset. To predict protein-designed cyclic peptide complex structure with AfDesign, we used alphafold2_multimer_v3 model (AF_v3). In this study, the input sequences were based on PDB files, If there is missing residues in the PDB, it is set to ‘X’. However, only sequence information was used as input for structure prediction, without using templates containing structural information. Among the resulting MSA files (`a3m` files), we prepared those where only the target protein sequence had MSA, and the peptide sequence had a single sequence, even if MSA was present in the peptide sequence. When predicting the complexes of peptide sequences designed in AfDesign and their target proteins, the same MSA was used without re-obtaining the MSA, and only the single peptide sequence-modified `a3m` files were used.

While localcolabfold was updated during our study, the commit history used was as follows: the localcolabfold repository, https://github.com/YoshitakaMo/localcolabfold/tree/e2bbc1aba7ecadbbea9265997cbda85bbcc4c75a (accessed on April 12, 2023) and ColabFold repository, https://github.com/sokrypton/ColabFold/commit/7d5e42c00829e12e38bd8965767302367cf90cd2 (accessed on May 3, 2023). The AF2 models used in localcolabfold were downloaded from the databases https://storage.googleapis.com/alphafold/alphafold_params_2022-03-02.tar and https://storage.googleapis.com/alphafold/alphafold_params_2022-12-06.tar (accessed on April 7, 2023).

### 2.2. AfDesign settings

We implemented a fixed-version cyclic peptide offset to the AfDesign binder hallucination protocol to design a cyclic peptide binder [55] (Figure 1**b**). Cyclic peptide binders for all proteins examined in this study were calculated using the AfDesign binder hallucination protocol [56]. All parameter settings were the same. To design a cyclic binder in AfDesign, the PDB file, which is the structural information of the target protein, and the length of the cyclic binder to be designed were set as inputs. In this study, only the structural information of the target protein was used, and the interface information of complexes such as hotspot was not used. For example, the MDM2 protein and p53 peptide complex, where the PDBID is 1YCR, chain A was used as the MDM2 structural template for AfDesign. The correspondence between other PDBIDs and the target protein chain is shown in Supplementary Table S1. Then, `binder_len` was set to 13, which was the same as the maximum length in a cyclic peptide design study using a cyclic offset [52]. The design method used was `design_pssm_semigreedy()`, where `soft_iter` and `hard_iter` were set to 120 and 32, respectively. Unless otherwise noted, the other settings were left at default. AfDesign was run 100 times for all target protein settings of the seeds from 1 to 100. The versions of `jax` and `jaxlib` used in this study were 0.3.25 and 0.3.25+cuda11.cudnn82, respectively. To ensure reproducibility, `TF_CUDNN_DETERMINISTIC=1` was set such that JAX behaved deterministically. While AfDesign was updated during our study, the commit history used was as follows: the ColabDesign repository—https://github.com/sokrypton/ColabDesign/tree/6d3837d94a63df561e73477dfdd553b0cf894a90 (accessed on April 7, 2023). The AF2 models used in AfDesign were downloaded from https://storage.googleapis.com/alphafold/alphafold_params_2022-12-06.tar (accessed April 7, 2023).

### 2.3. Calculation of protein-peptide local docking for cyclic peptide

The target protein and designed peptide were calculated using ADCP which is a state-of-the-art cyclic peptide docking tool within the AutoDockFR software suite version 1.0 [37,57]. This procedure is described in the AutoDockFR document. First, the PDB files prepared as receptors and peptides were protonated using reduce version 3.23.130521 [58], and then the protonated PDB files were converted to PDBQT files using two scripts, `prepare_ligand` and `prepare_receptor`, of the AutoDockFR software suite. In addition, a docking box was placed and sized over the receptor pocket into which the peptide was docked, and a `trg` file was calculated using `agfr` to compute an affinity map for the list of atom types in AutoDock 4 [59]. Up to 2.5 million evaluations of the scoring function were performed to increase the likelihood of finding the best docking pose (the global minimum of the scoring function), and 50 independent searches were performed for each. The ADCP argument was `-N 50 -n 2500000`. The “`-cyc`” option was used when predicting cyclic peptides.

### 2.4. Visualization of sequence logos

The sequence with the lowest final loss value in AfDesign binder hallucination protocol in the latter 32 iterations (`hard_iter`) in `design_pssm_semigreedy()` was selected as the best sequence. The sequence logos were created using the WebLogo 3 Python package (version 3.7.9) [60]. The designed cyclic peptide sequences were used as inputs and aligned in PyMOL considering the 3D coordinates of the native peptide (Supplementary Figure S1).

1. The PDB target protein was aligned with target proteins in the complex with cyclic peptides predicted by AF2.
2. A window with each native peptide and cyclic peptide shifted by one residue was created with a width of 6 residues (from the first residue to the sixth residue, from the second residue to the seventh residue and so on. The native peptide windows will have a length of −5, and the cyclic peptide will have the same number of windows as its length).
3. The RMSD of the C*α* of each window in 1 was calculated using rms_cur. Using 1YCR as an example, (13−5) × 13 = 104 RMSDs were calculated per design.
4. To compare with native “linear” peptides, using the window index with the lowest RMSD of the C*α* among them, the design sequence was aligned based on the index numbers of the native peptide sequence and the cyclic peptide. The lowest RMSD was named RMSD_best in this study.
5. After alignment, the non-6 letters were filled in with ‘-’. Sequence logos were created via WebLogo using the thresholding alignment sequence in RMSD_best as input (Figure 1**c**). The sequence logo represents each column of alignment as a stack of letters, with the height of each letter proportional to the observed frequency of the corresponding amino acid, and the overall height of each stack proportional to the degree of sequence conservation (measured in bits) at that location. The maximum sequence conservation per site is log_2_ 20 ≃ 4.32 bits for amino acids. The width of the letters is proportional to ungaps of each column (the more ‘-’ there are in each column instead of amino acid represents, the thinner the letters).

If the native peptide was smaller than six residues, its length was adjusted to the length of the native peptide. For linear peptides, the window width was set to 10 residues.

### 2.5. Rosetta interface analyzer

The Rosetta interface scores (dG_separated/dSASA × 100) for the predicted protein and cyclic peptide structural models were calculated using a Rosetta interface analyzer [61]. First, “minimize” was used to obtain the minimum energy structure around the initial predicted complex structure. To prevent cyclic peptides in the complex from becoming linear peptides, we used the “`-use_truncated_termini`” and “`-relax:bb_move false`” options and the minimized complex as the input “InterfaceAnalyzer.” To repack the exposed interfaces when calculating binding energy, we used the option “`-pack_separated`”. Standard weights, REF15, and Rosetta 3.13 Linux Release were used for all Rosetta applications in this study.

### 2.6. Calculation of solubility and lipophilicity

As a preprocessing step, SMILES were prepared for linear peptides through acetylation of the N-terminus of the peptide sequence and amidation of the C-terminus using RDKit `Chem.MolFromHELM()`, and conformers were prepared using LigPrep in the Schrödinger software suite (version 2021-1). Finally, QPLogS and QPLogPo/w values were calculated for solubility and lipophilicity using QikProp in the Schrödinger software suite (version 2021-1), respectively. If multiple conformers were generated from a single peptide sequence, the logS and logP values were averaged for each peptide sequence.

### 2.7. Interatomic interactions between PD-L1 and the designed cyclic peptide

The online tool PLIP [62] (https://plip-tool.biotec.tu-dresden.de, accessed on July 3, 2023) was used to analyze the atomic interactions between the designed peptide and PD-L1. To detect these interactions, the inter/peptide mode was selected, and the chain ID of the peptide was entered in the advanced options. The structure of the designed sequence was the top-ranked structure predicted by localcolabfold. The predicted structure of the designed sequence with the lowest Rosetta interface analyzer score was greater than predicted local distance difference test (pLDDT) by over 90.

## 3. Results

### 3.1. Structure prediction of the protein and cyclic peptide complex

Before designing cyclic peptides that target proteins in protein-peptide complexes, an offset matrix of relative positions must be encoded specifically for protein-cyclic peptide complexes to allow structure prediction with AF2. We developed a cyclic peptide complex offset based on the offset for single cyclic peptides used in a previous study [52]. Recently, a fixed cyclic offset implementation was reported and applied to a cyclic peptide complex offset [55]. The part of ColabFold encoding relative position was modified (See Materials and Methods). As shown in Figure 1**a**, the offset matrix for the relative position encoding the protein-cyclic peptide complex was designed such that the offset in the protein region was the same as the default offset, whereas the offset in the peptide region was cyclic.

To evaluate how accurately AF2 with the developed cyclic peptide complex offset could predict protein-cyclic peptide complexes, we selected five complexes that were also used to validate the cyclic peptide docking tool ADCP [57]. Among the three head-to-tail types reported by the ADCP, the following were selected: one complex from the SFTI-1 cyclic peptide complex group (1SFI), two complexes from the HIV integrase complex with the cyclic peptide group (3AV9 and 3WNE), and two complexes from another group (3ZGC and 5XN3). In the latest version of the AF2_v3 training set, only peptides with fewer than four amino acid residues were excluded, including most cyclic peptide complexes registered in the PDB. However, there were no training sets for the alphafold2_ptm model (AF2_ptm) and AF2_v2, because the training excluded peptides with fewer than 16 amino acid residues [52,63]. The AF2_ptm was developed to predict single proteins [39]. This model can predict complexes by using glycine linkers and chain breaks [44,45,54]. On the other hand, AF2_v2 was especially developed to predict protein complexes, achieving higher accuracy than AF2_ptm by using a glycine linker and chain break [42]. It was recently reported that AF2 with AF2_ptm could be used to predict cyclic peptides using cyclic offsets with high accuracy. We compared the results for AF2_ptm and AF2_v2 using a cyclic peptide complex offset. AF2_v2 showed higher pLDDT and predicted aligned error (PAE) values than alphafold2_ptm (Supplementary Figure S2). Therefore, AF2_v2 was considered suitable for predicting protein-cyclic peptide complexes. Next, we compared the prediction accuracy of AF2_v2 using the cyclic peptide complex offset to that of ADCP [57]. Five protein-cyclic peptide complex structures predicted using AF2_v2 with the cyclic peptide complex offset and structure with lowest “ref.rmsd” among the top 10 ADCP local docking are shown in Figure 2, and the RMSD results after alignment with native target proteins and target peptide are shown in Table 1. In complex prediction by AF2_v2 using the cyclic peptide complex offset, alignment with the native and predicted structure target proteins and calculation of the cyclic peptide RMSD resulted in a high accuracy of 0.87 Å to 4.28 Å. In contrast, the RMSD of ADCP ranged from 3.40 Å to 7.09 Å, which was higher than that of the AF2_v2 complex prediction using the cyclic peptide complex offset, which also showed a higher RMSD, except for 5XN3. However, while ADCP performed local docking using information on the binding sites between native cyclic peptides and the target protein as the docking grid, AF2_v2 involved a completely blind prediction, because these cyclic peptides are not included in the training dataset. Therefore, accuracy comparison is disadvantageous for AF2. The accuracy of predicting the structure as a cyclic peptide was evaluated in the same manner. In all cases, AF2_v2 had a lower RMSD than ADCP (Table 1). This shows that the 5XN3 cyclic peptide could be predicted more accurately than ADCP as a single peptide. However, it was difficult to predict the appropriate position for the complex structure. These results suggest that protein-cyclic peptide complex prediction with AF2_v2 using the cyclic peptide complex offset can be performed with equivalent or better accuracy than that of ADCP in many cases.

**Table 1.** Comparison of cyclic peptide RMSD for the protein-cyclic peptide complex structures predicted by AF2_v2 and ADCP (based on align native protein and align native peptide)

|  |  | 1SFI | 3AV9 | 3WNE | 3ZGC | 5XN3 |
| --- | --- | --- | --- | --- | --- | --- |
| RMSD (Å)<br>(Aligned with protein) | AF2_v2 | 0.86 | 1.57 | 1.63 | 1.85 | 4.28 |
|  | ADCP | 6.71 | 7.09 | 4.29 | 3.58 | 3.40 |
| RMSD (Å)<br>(Aligned with peptide) | AF2_v2 | 0.53 | 1.01 | 1.37 | 1.23 | 2.94 |
|  | ADCP | 5.01 | 3.32 | 2.49 | 2.27 | 3.28 |

**Figure 2.**
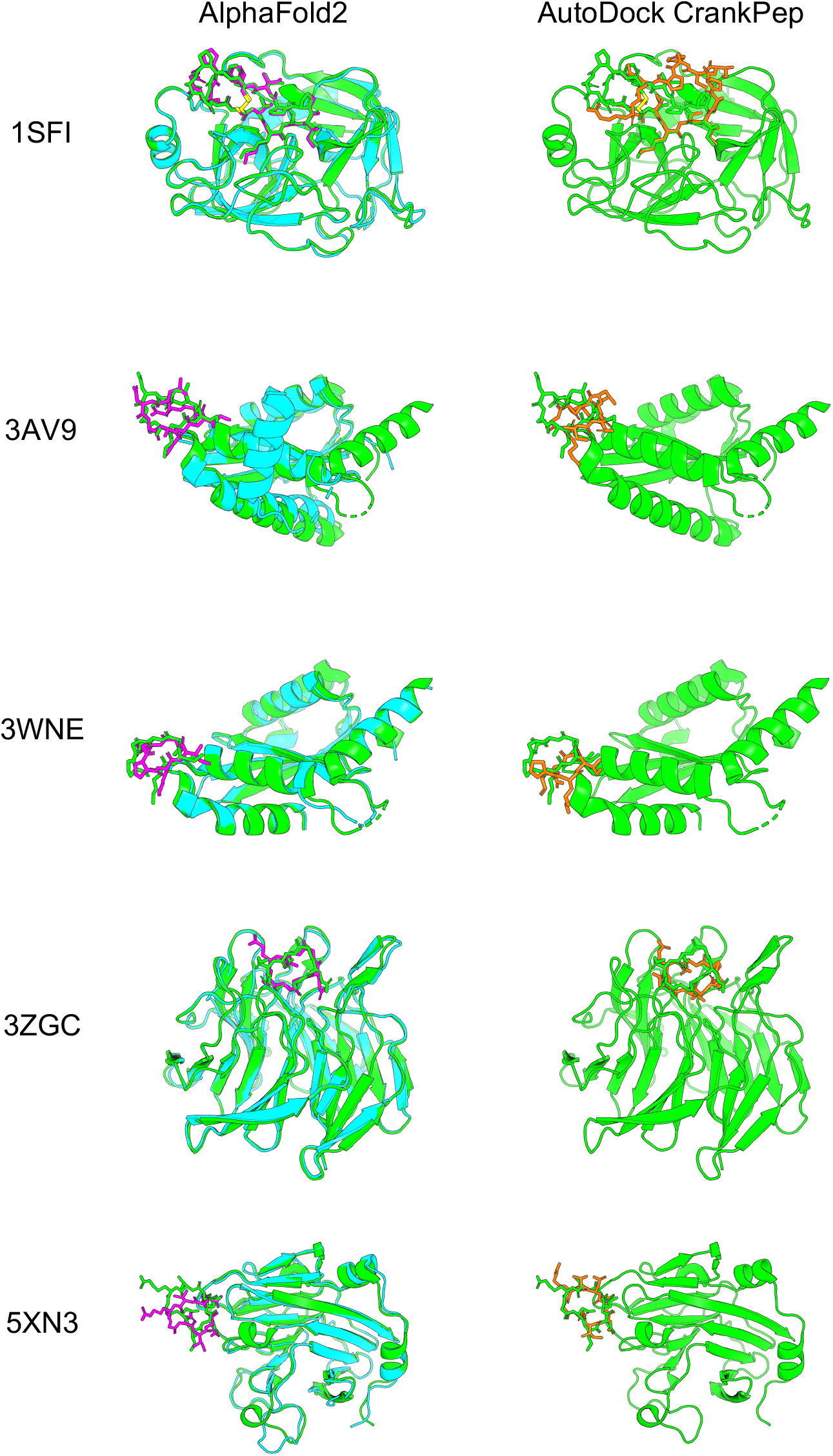
Comparison of predicted protein-cyclic peptide complex structures of AF2 and ADCP. The left column is the predicted structure of AF2_v2 with the cyclic peptide complex offset. The right column is ADCP local docking structure with the lowest “ref.rmsd” among the top 10 structures. Green shows the predicted structure of the PDB target protein, cyan shows the predicted structure of the AF2 target protein, and magenta shows the predicted structure of the AF2 cyclic peptide. Orange shows the ADCP local docking cyclic peptide structure.

### 3.2. Cyclic peptide binder hallucination targeting the protein-peptide complex

AfDesign is a public module also known as ColabDesign with AF2 developed by Dr. Sergey Ovchinnikov [56]. AF2 is used as a structure prediction oracle; the optimization target is the binding sequence, and the loss function is defined as a flexible function of AF2 confidences. It outputs the pLDDT, PAE, and distogram, among which pLDDT measures the quality of the local model for each residue, PAE indicates the confidence level of each amino acid residue pair, and distogram indicates the distance prediction probability of C_*β*_ for an amino acid residue pair. We used AfDesign to prepare a solubility-aware peptide binder [51]. A previous study was motivated by the binder design using AfDesign binder hallucinations, which had just been released as an open source, which tended to yield many binders with low solubility. Therefore, we added a solubility index to the loss function to improve the solubility of the designed binders. AfDesign has undergone several updates and feature additions by Dr. Sergey Ovchinnikov. Therefore, to begin this study, we compared the degree of difference in solubility and lipophilicity between versions 1.1.1 and beta. V1.1.1 solved the solubility and lipophilicity problems better than the default beta version, and the QPLogS and QPLogPo/w distributions were almost the same as when the hydropathy index was used as the solubility index (Supplementary Figure S3). Owing to improvements in the optimizer and loss function weights, the AfDesign binder hallucination protocol in v1.1.1 reduced the solubility problem, compared to the beta version. Therefore, we used v1.1.1 in this study. Figure 1 **b** shows our approach to protein-bound cyclic peptide design, which was performed using the AfDesign binder hallucination protocol with our cyclic peptide complex offset. In the design, we did not need to consider the scope of the AF2 training set. Therefore, we evaluated which was more desirable for cyclic peptide binder hallucination using AfDesign: AF2_ptm, or AF2_v3. To evaluate the quality of the designed sequences, we used pLDDT of the cyclic peptide sequence in its predicted protein complex. As shown in Supplementary Figure S4, the design sequence of AF2_ptm tended to have a higher pLDDT than AF2_v3 for 1YCR, indicating that AF2_ptm is a desirable model for cyclic peptide binder hallucinations in AfDesign.

Next, we verified whether the high pLDDT cyclic binder could be hallucinated against MDM2 as the target protein. The structure predicted by ColabFold for the designed sequences was scored using the Rosetta interface analyzer. It approximates the binding energy by calculating the change in Rosetta energy when the separated binding partners come together to form a complex with no change between the bound and unbound monomer structures [61]. However, the predicted structure of AF2 is often not the minimum of the Rosetta energy function, and it often exhibits higher energy values. Therefore, the relative energies cannot be compared directly. To solve this problem, we performed the Rosetta minimization protocol on the AF2 predicted structures, as seen in previous studies [64]. This protocol is based on a simple gradient to determine the minimum energy structure around the initial structure. The Rosetta interface analyzer was used after converging the AF2 predicted structure to the closest local minimum under the Rosetta energy function, while maximally guaranteeing the original AF2 predicted structure.

We compared the scores of the designed structure and target protein with those of the native structure, p53 peptide, and MDM2 using scores normalized to solvent accessible surface area (SASA). As shown in Figure 3**a** and **b**, the SASA of all design sequence structures was smaller than that of the native structure; however, 88% of the design sequence binding energies were lower than that of the native structure (lower binding energy is more stable), and 79.5% had a pLDDT over 70. This means that the contact areas of the peptide with the designed sequences were smaller than those of the native peptide, but the former was more favorable in terms of surface area per unit. The predicted design structures were filtered to select those with binding energies higher than the native structure and similar peptide interface structures. The structure with pLDDT over 70, binding energy lower than that of the native structure, and the lowest RMSD_best, which considers the 3D alignment of cyclic and native peptides (see Materials and Methods), was chosen among the predicted design structures. The results are shown in Figure 3 **c**, with a pLDDT of 91.0, an interface analyzer score of − 2.456, and an RMSD_best of 0.718 Å. The same aromatic amino acid, phenylalanine, was placed at the 3rd phenylalanine and 7th tryptophan positions in the native peptide sequence that interacts with the MDM2 hotspot in the predicted design structure.

**Figure 3.**
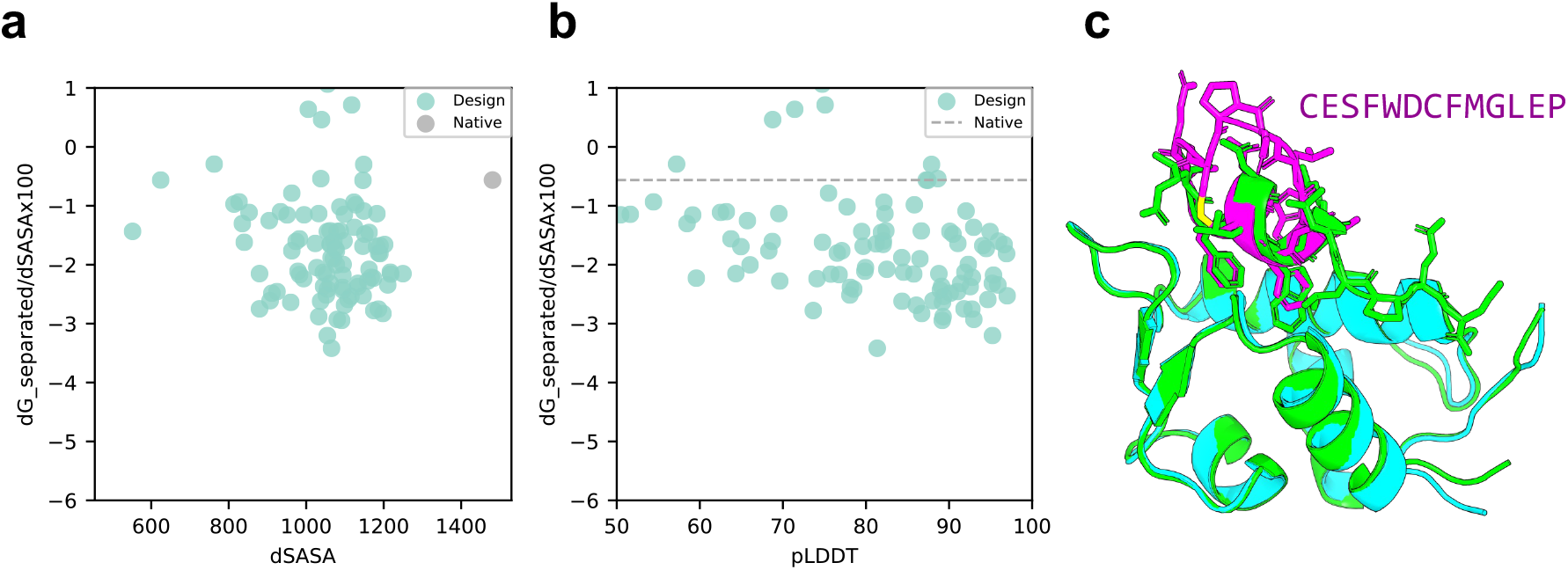
Comparison of binding energy using the Rosetta interface analyzer score between the native structure and the MDM2-targeted design sequence predicted structure. (**a**) Scatterplot of binding energy and SASA from the Rosetta interface analyzer. The grey dot presents the native structure value. (**b**) Scatterplot of binding energy from the Rosetta interface analyzer and pLDDT. The grey line presents the native structure score. (**c**) Comparison of crystal 1YCR structure and AfDesign sequence structure predicted using ColabFold. The predicted structure of the design sequence had a Rosetta interface analyzer score lower than that of the native structure and had the lowest RMSD_best. Green shows the native target protein structure, cyan shows the predicted target protein structure, and magenta shows the predicted cyclic peptide structure.

Next, a sequence logo was created through proper alignment of the designed and native sequences (see Materials and Methods and Supplementary Figure S1). The designed sequences of the cyclic and linear peptides were compared to determine the effect of cyclic peptide complex offset. As shown in Figure 4, the sequence logos of cyclic and linear peptide design sequences were created using RMSD_best for native peptides as the threshold. The designed cyclic peptide sequences tended to be hallucinated mainly in the 3rd to 8th native positions, with the 3rd and 7th residues interacting with the MDM2 hotspot region, consisting mainly of phenylalanine and tryptophan. This was similar to the designed sequence logos of the linear peptides. This result indicates that the cyclic peptide complex offset had little effect on critical residues at the binding interface, allowing for cyclic design.

**Figure 4.**
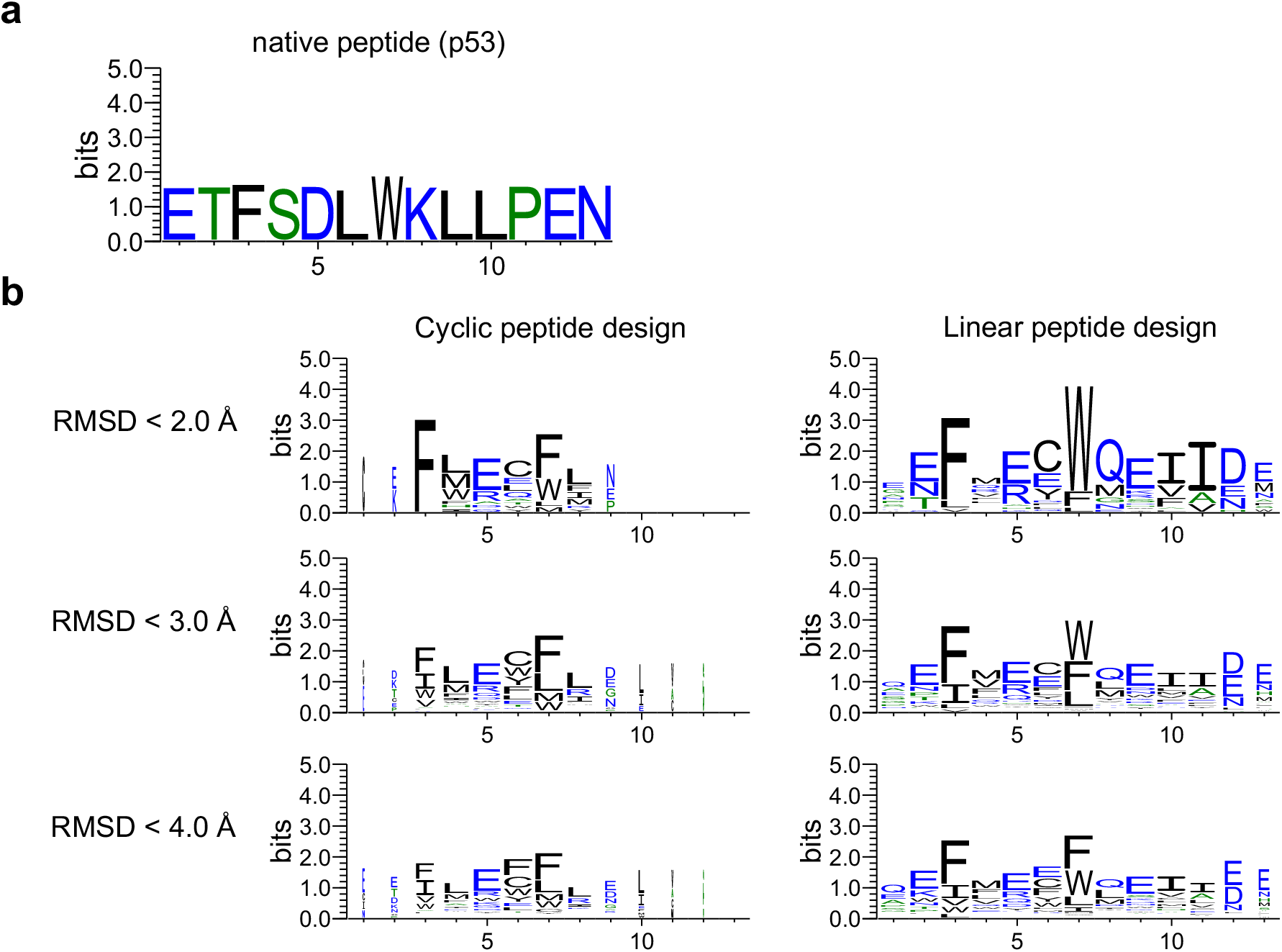
Comparison of sequence logos for cyclic and linear peptide designs. The sequence logo represents each column of alignment as a stack of letters, with the height of each letter proportional to the observed frequency of the corresponding amino acid, and the overall height of each stack proportional to the degree of sequence conservation at that location. The width of the letters is proportional to ungaps of each column. (**a**) Native peptide (p53) sequence logo. (**b**) The left column shows the sequence logo of cyclic design sequences for 1YCR with each RMSD_best threshold. The right column shows the linear design sequences for 1YCR with each RMSD_best threshold. Hydrophilic amino acids (R, K, D, E, N, and Q) are shown in blue, neutral amino acids (S, G, H, T, A, and P) are shown in green, and hydrophobic amino acids (Y, V, M, C, L, F, I, and W) are shown in black.

## 4. Discussion

### 4.1. Application of cyclic peptide binder hallucinations to other target protein-peptide complexes

To confirm whether cyclic peptide binder hallucinations could be applied to proteins other than 1YCR, we also applied them to target proteins of other protein–peptide complexes. The protein-peptide complex dataset was selected based on two previous studies for this evaluation, using 12 protein-peptide complexes as target proteins, for which AF can be predicted within 2 Å RMSD [45,65]. The same processes were used to design cyclic peptide binders for the 12 protein-peptide complexes. The predicted design structures of cyclic peptides for the target proteins of each protein-peptide complex were filtered in the same manner as 1YCR. However, if no structure had a binding energy lower than that of native PDB, the binding energy was set to less than 0. Figure 5 shows the predicted designs resulting from these filters and the structures aligned with the native target proteins. Table 2 lists the pLDDT for the cyclic peptides, Rosetta interface analyzer scores, and RMSD_best of the predicted design structures.

**Table 2.**
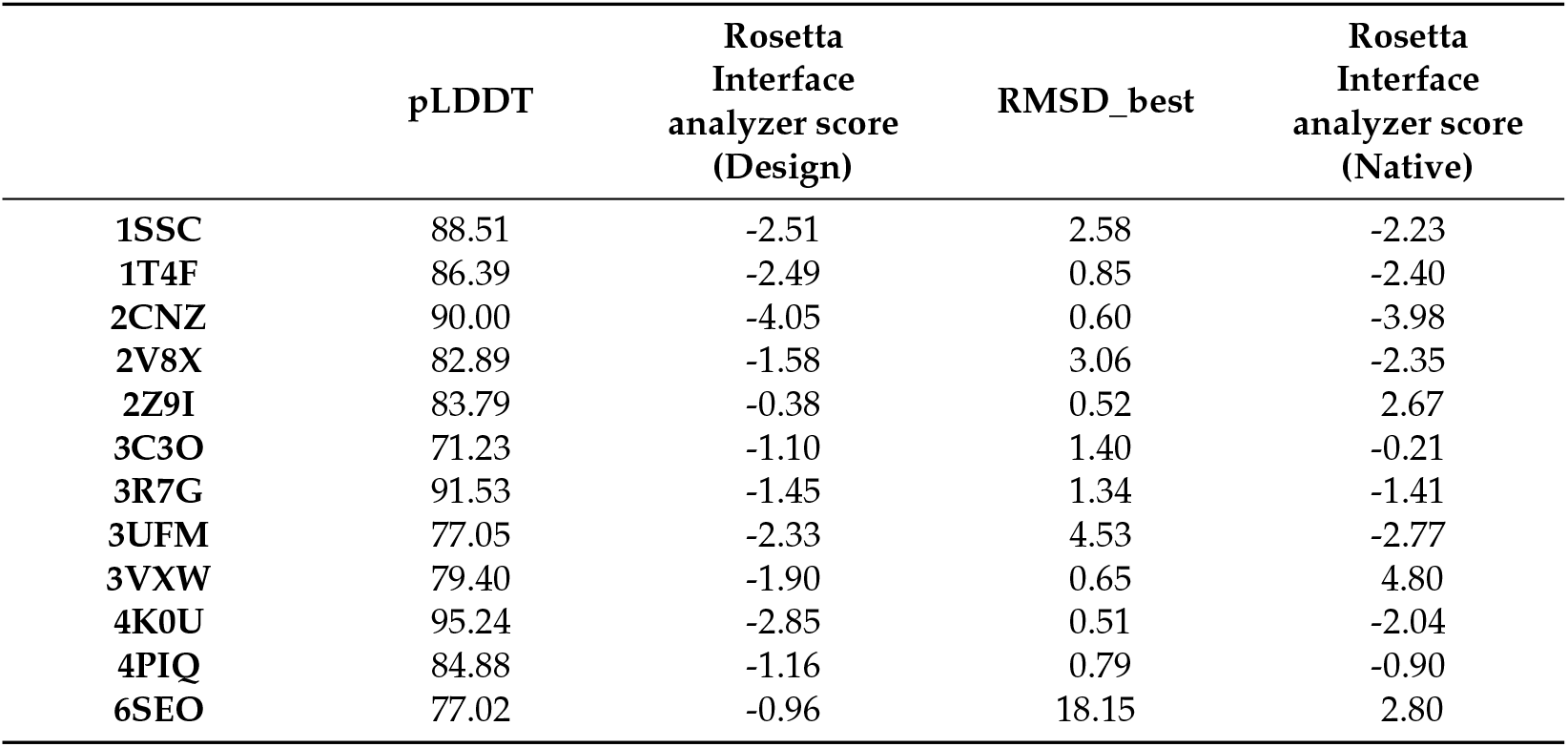
pLDDT, Rosetta interface analyzer score, and RMSD_best of the structures of cyclic peptide binders for 12 protein-cyclic peptide complexes predicted using AF2_v3.

|  | pLDDT | Rosetta<br>Interface<br>analyzer score<br>(Design) | RMSD_best | Rosetta<br>Interface<br>analyzer score<br>(Native) |
| --- | --- | --- | --- | --- |
| 1SSC | 88.51 | -2.51 | 2.58 | -2.23 |
| 1T4F | 86.39 | -2.49 | 0.85 | -2.40 |
| 2CNZ | 90.00 | -4.05 | 0.60 | -3.98 |
| 2V8X | 82.89 | -1.58 | 3.06 | -2.35 |
| 2Z9I | 83.79 | -0.38 | 0.52 | 2.67 |
| 3C3O | 71.23 | -1.10 | 1.40 | -0.21 |
| 3R7G | 91.53 | -1.45 | 1.34 | -1.41 |
| 3UFM | 77.05 | -2.33 | 4.53 | -2.77 |
| 3VXW | 79.40 | -1.90 | 0.65 | 4.80 |
| 4K0U | 95.24 | -2.85 | 0.51 | -2.04 |
| 4PIQ | 84.88 | -1.16 | 0.79 | -0.90 |
| 6SEO | 77.02 | -0.96 | 18.15 | 2.80 |

**Figure 5.**
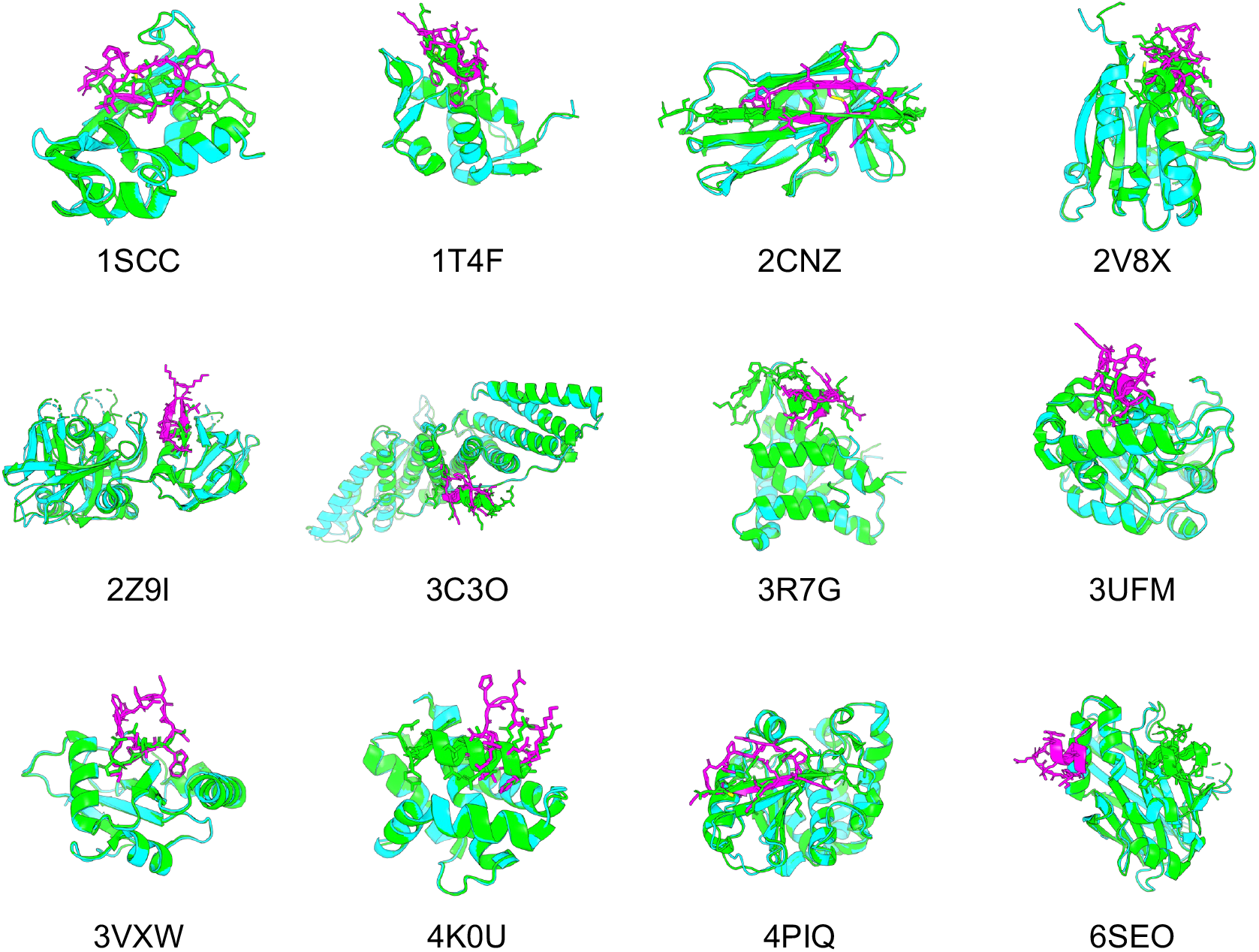
Comparison of the predicted structures of 12 target proteins and designed cyclic peptides with the native PDB structures. Green shows the predicted structure of the PDB target protein, cyan shows the predicted structure of the AF2 target protein, and magenta shows the predicted structure of the AF2 cyclic peptide.

For nine of the 12 structures, the Rosetta interface analyzer score was lower for the predicted design than for the native structure and RMSD_best was lower than 3 Å, with a mean pLDDT for the predicted cyclic peptide design of 85.66 and a mean RMSD_best of 1.03 Å, suggesting its high conference and similarity to the native peptide structures. The predicted structures of each design were as follows: 1SSC had a short *β*-sheet structure and disulfide bonds, 1T4F had matching phenylalanine and tryptophan residues known to interact with the hotspot of the target protein, and RMSD_best was 0.85 Å. Further, 2CNZ had two *β*-sheet structures and disulfide bonds, and the arginine and threonine atoms of the design matched well with the positions of the native glutamine and threonine. There were two *β*-sheet structures in 2Z9I. A helical structure was seen in 3C3O, and the tryptophan position was consistent with its native structure. The helical structure in 3R7G was similar to that of the C-terminal side in the native protein. The tryptophan and isoleucine in 3VXW were the same as the native ones, and their backbones matched well from tryptophan to isoleucine. In addition, 3VXW had the same tryptophan and isoleucine structures as its native form, whereas 4K0U had a helical structure similar to that of the native form, and its tryptophan position was well matched. Finally, 4PIQ had a single *β*-sheet structure, and its cysteine was identical to that of the native protein. In its native form, this cysteine forms a disulfide bond to the target protein, which is considered a very important site. Overall, 74% of the 4PIQ designs had a cysteine in this position.

However, these three complexes were not well designed. Specifically, 2V8X and 3UFM had no design with a Rosetta interface analyzer score lower than that of the native structure, and their RMSD_ best values were higher than those of the other structures. In addition, 6SEO had an RMSD_best of 18.15 Å, and the predicted structure was clearly in a different location from that of the native peptide. The RMSD_best distribution of the predicted design structures for the 12 complexes indicated that these three complexes were less likely to be hallucinated at the native peptide position via cyclic peptide binder hallucinations (Supplementary Figure S5).

### 4.2. Cyclic peptide binder hallucination targeting protein and protein complex

The focus of this study was to design cyclic peptide binders for protein-peptide complexes. As a challenge, we designed cyclic peptide binders for protein-protein complexes. We evaluated PD-L1 as a target protein of the PD-1/PD-L1 complex (PDB ID: 4ZQK), which is involved in the cancer cell cycle checkpoints [66,67]. This is challenging in terms of designing binders for broad *beta*-sheet interactions, unlike in protein-peptide complexes. As shown in Figure 6**a** and **b**, the SASA of the designed cyclic peptide was approximately 42% that of the native SASA; however, 44% had a normalized binding energy lower than that of the native SASA. Of these, 22.7% (10 of 44) had a pLDDT of over 90 (i.e., 10% of the total). The predicted structure of the design sequence with the lowest Rosetta interface analyzer score and pLDDT over 90 is shown in Figure 6**c**. The predicted structure of the designed sequence showed that the helical and loop structures of the cyclic peptide formed in the *β*-sheet and loop structures of PD-1, respectively, at the interaction site. Additionally, PLIP was used to analyze the atomic interactions between the designed cyclic peptide and PD-L1 [62]. There were hydrogen bonds between Glu58 and Tyr123 in the target protein and Tyr1 and Asp8 in the cyclic peptide, a salt bridge between the target protein Glu58 and the cyclic peptide Arg13. Hydrophobic interactions were also observed between Arg113,

**Figure 6.**
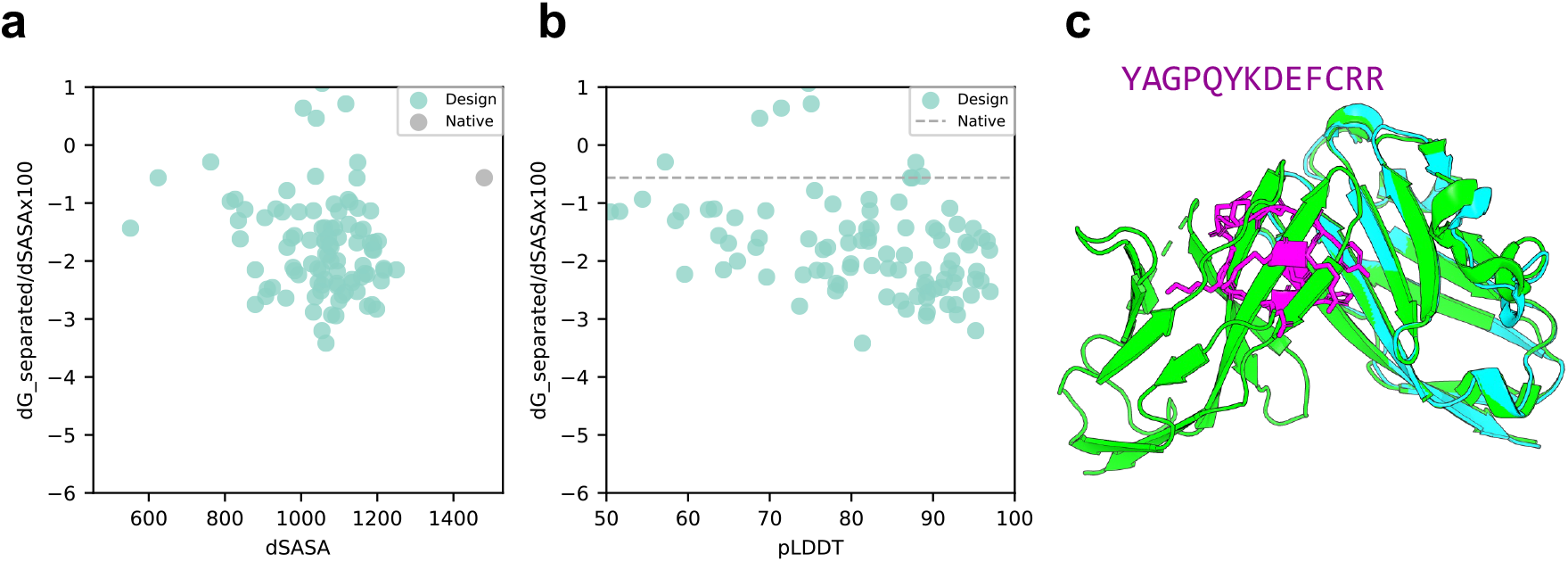
Comparison of binding energy using the Rosetta interface analyzer score between the native structure and PD-L1-targeted design predicted structure. (**a**) Scatter plot of binding energy and SASA from the Rosetta interface analyzer. The grey dot shows the native structure value. (**b**) Scatterplot of binding energy from the Rosetta interface analyzer and pLDDT. The grey line shows the native structure score. (**c**) Comparison of crystal 4ZQK structure and AfDesign sequence structure predicted via ColabFold. The predicted structure with the lowest Rosetta interface analyzer score and pLDDT over 90 is shown. Green shows the native target protein structure, cyan shows the predicted target protein structure, and magenta shows the predicted cyclic peptide structure.

Met115, and Tyr123 of the target protein and Phe10 of the cyclic peptide, Tyr56 of the target protein, and Tyr1 of the cyclic peptide, respectively (Supplementary Figure S6).

### 4.3. Limitations and challenges

Using the cyclic peptide complex offset developed in this study, we tested whether protein and cyclic peptide complex prediction using AF2 models could be used in the same input method as conventional AF2 complex prediction. Comparing its accuracy with other methods, such as ADCP and other AF2 models, using larger datasets will be a challenge for future work.

Binder hallucinations using AfDesign can hallucinate desired structures with a flexible loss function. However, some target proteins may be hallucinated at a location clearly different from that of the native peptide, with which they interact (Supplementary Figure S5). Possible solutions include increasing the number of designs or specifying hotspots to increase the possibility of hallucinations in a relatively more appropriate location. However, this remains a challenge for the future.

## 5. Conclusions

We developed a specific protein-cyclic peptide complex offset and implemented it in ColabFold to predict the RMSD of the cyclic peptide region in head-to-tail cyclic peptide complex prediction with a high accuracy of 0.86 Å to 1.85 Å, except for one example. This outstanding achievement exceeded the accuracy of ADCP, a state-of-the-art cyclic peptide-docking tool. Furthermore, applying AfDesign peptide binder hallucination to the cyclic peptide complex offset enabled de novo binder hallucination of cyclic peptides, with MDM2 as the target protein. A total of 88% of the designed sequences had lower SASA-normalized binding energies than the native peptides. Furthermore, 27% of all sequences had pLDDT > 90. Furthermore, the alignment of the designed sequences and drawing of sequence logos showed that the positions of the aromatic amino acids important for binding to the native peptides were consistent. We also designed cyclic peptides for the 12 target proteins of the protein-peptide complexes selected in a previous study. Nine of the 12 sequences were designed with a lower normalized binding energy than the native structure and RMSD_best was lower than 3 Å, their mean pLDDT was 85.66, and the mean RMSD_best was 1.03 Å. We also designed cyclic peptides targeting PD-L1 in the PD-1/PD-L1 complex and found that 10% of the sequences had a pLDDT > 90 and a binding energy lower than the normalized binding energy of the native structure. In this study, we showed that our method could be used to design putative cyclic peptides targeting PPI using AfDesign binder hallucinations with cyclic peptide complex offsets for proteins and cyclic peptide complexes.

## Author Contributions

Conceptualization, T.K. and M.O.; methodology, T.K. and M.O.; software, T.K.; validation, T.K.; formal analysis, T.K. and M.O.; investigation, T.K. and M.O.; resources, T.K.; data curation, T.K.; writing—original draft preparation, T.K. and M.O; writing—review and editing, T.K. and M.O; visualization, T.K.; supervision, M.O.; project administration, M.O.; funding acquisition, M.O. All authors have read and agreed to the published version of the manuscript.

## Funding

This work was financially supported by the Japan Science and Technology Agency (JST) FOREST (Grant No. JPMJFR216J) and JST ACT-X (Grant No. JPMJAX20A3), Japan Society for the Promotion of Science KAKENHI (Grant Nos. 23H03496, 23H04880, and 23H04887), and Japan Agency for Medical Research and Development Basis for Supporting Innovative Drug Discovery and Life Science Research (Grant No. JP22ama121026).

## Institutional Review Board Statement

Not applicable.

## Informed Consent Statement

Not applicable.

## Data Availability Statement

The implementation and experimental data are available on an open-source basis at https://github.com/ohuelab/ColabFold-cycpep-dock (accessed on 17 July 2023). and https://github.com/ohuelab/ColabDesign-cyclic-binder (accessed on 17 July 2023).

## Acknowledgments

The computational experiments were performed using the TSUBAME 3.0 super-computer at the Tokyo Institute of Technology. Sergey Ovchinnikov, John Jumper, and DeepMind AlphaFold team informed us of the amino acid length cutoff in the AlphaFold2 training dataset. Naoya Kobayashi and Shingo Honda advised us on how to minimize protein-cyclic peptide complexes.

## Conflicts of Interest

The authors declare no conflicts of interest.

## Abbreviations

The following abbreviations are used in this manuscript:

PPI: protein-protein interaction
HTVS: High-throughput virtual screening
DL: deep learning
AF2: AlphaFold2
RMSD: root mean square deviation
MSA: multiple sequence alignment
PDB: Protein Data Bank
ADCP: AutoDock CrankPep
PAE: predicted aligned error
pLDDT: predicted local distance difference test
AF2_v3: alphafold2_multimer_v3 model
AF2_v2: alphafold2_multimer_v2 model
AF2_ptm: alphafold2_ptm model
SASA: solvent accessible surface area

## Notes

### Competing Interest Statement

The authors have declared no competing interest.

## References

1. Bonetta, L. Interactome under Construction. Nature 2010, 468, 851–852. https://doi.org/10.1038/468851a.

2. Oughtred, R.; Rust, J.; Chang, C.; Breitkreutz, B.-J.; Stark, C.; Willems, A.; Boucher, L.; Leung, G.; Kolas, N.; Zhang, F.; et al. The BioGRID Database: A Comprehensive Biomedical Resource of Curated Protein, Genetic, and Chemical Interactions. Protein Sci. 2021, 30, 187–200. https://doi.org/10.1002/pro.3978.

3. Toogood, P.L. Inhibition of Protein-Protein Association by Small Molecules: Approaches and Progress. J. Med. Chem. 2002, 45, 1543–1558. https://doi.org/10.1021/jm010468s.

4. Arkin, M.R.; Wells, J.A. Small-Molecule Inhibitors of Protein–Protein Interactions: Progressing towards the dream. Nat. Rev. Drug Discov. 2004, 3, 301–317. https://doi.org/10.1038/nrd1343.

5. Dev, K.K. Making Protein Interactions Druggable: Targeting PDZ Domains. Nat. Rev. Drug Discov. 2004, 3, 1047–1056. https://doi.org/10.1038/nrd1578.

6. Jin, L.; Wang, W.; Fang, G. Targeting Protein-Protein Interaction by Small Molecules. Annu. Rev. Pharmacol. Toxicol. 2014, 54, 435–456. https://doi.org/10.1146/annurev-pharmtox-011613-140028.

7. Ivanov, A.A.; Khuri, F.R.; Fu, H. Targeting Protein–Protein Interactions as an Anticancer Strategy Trends Pharmacol. Sci. 2013, 34, 393–400. https://doi.org/10.1016/j.tips.2013.04.007.

8. Shin, W.-H.; Kumazawa, K.; Imai, K.; Hirokawa, T.; Kihara, D. Current Challenges and Opportunities in Designing Protein–Protein Interaction Targeted Drugs. Adv. Appl. Bioinform. Chem. 2020, 13, 11–25.

9. Kosugi, T.; Ohue, M. Quantitative Estimate Index for Early-Stage Screening of Compounds Targeting Protein-Protein Interactions. Int. J. Mol. Sci. 2021, 22, 10925. https://doi.org/10.3390/ijms222010925.

10. Vinogradov, A.A.; Yin, Y.; Suga, H. Macrocyclic Peptides as Drug Candidates: Recent Progress and Remaining Challenges. J. Am. Chem. Soc. 2019, 141, 4167–4181, https://doi.org/10.1021/jacs.8b13178.

11. Muttenthaler, M.; King, G.F.; Adams, D.J.; Alewood, P.F. Trends in Peptide Drug Discovery. Nat. Rev. Drug Discov. 2021, 20, 309–325, https://doi.org/10.1038/s41573-020-00135-8.

12. Tsomaia, N. Peptide Therapeutics: Targeting the Undruggable Space. Eur. J. Med. Chem. 2015, 94, 459–470, https://doi.org/10.1016/j.ejmech.2015.01.014.

13. Whitty, A.; Zhong, M.; Viarengo, L.; Beglov, D.; Hall, D.R.; Vajda, S. Quantifying the Chameleonic Properties of Macrocycles and Other High-Molecular-Weight Drugs. Drug Discov. Today 2016, 21, 712–717, https://doi.org/10.1016/j.drudis.2016.02.005.

14. Lee, D.; Lee, S.; Choi, J.; Song, Y.-K.; Kim, M.J.; Shin, D.-S.; Bae, M.A.; Kim, Y.-C.; Park, C.-J.; Lee, K.-R.; et al. Interplay among Conformation, Intramolecular Hydrogen Bonds, and Chameleonicity in the Membrane Permeability and Cyclophilin A Binding of Macrocyclic Peptide Cyclosporin O Derivatives. J. Med. Chem. 2021, 64, 8272–8286, https://doi.org/10.1021/acs.jmedchem.1c00211.

15. Sugita, M.; Sugiyama, S.; Fujie, T.; Yoshikawa, Y.; Yanagisawa, K.; Ohue, M.; Akiyama, Y. Large-scale membrane permeability prediction of cyclic peptides crossing a lipid bilayer based on enhanced sampling molecular dynamics simulations. J. Chem. Inf. Model. 2021, 61(7), 3681–3695, https://doi.org/10.1021/acs.jcim.1c00380

16. Sugita, M.; Fujie, T.; Yanagisawa, K.; Ohue, M.; Akiyama, Y. Lipid composition is critical for accurate membrane permeability prediction of cyclic peptides by molecular dynamics simulations. J. Chem. Inf. Model. 2022, 62(18), 4549–4560, https://doi.org/10.1021/acs.jcim.2c00931

17. Wu, C.-H.; Liu, I.-J.; Lu, R.-M.; Wu, H.-C. Advancement and Applications of Peptide Phage Display Technology in Biomedical Science. J. Biomed. Sci. 2016, 23, 8, https://doi.org/10.1186/s12929-016-0223-x.

18. Goto, Y.; Suga, H. The RaPID Platform for the Discovery of Pseudo-Natural Macrocyclic Peptides. Acc. Chem. Res. 2021, 54, 3604–3617, https://doi.org/10.1021/acs.accounts.1c00391.

19. Yamaguchi, J.; Naimuddin, M.; Biyani, M.; Sasaki, T.; Machida, M.; Kubo, T.; Funatsu, T.; Husimi, Y.; Nemoto, N. CDNA Display: A Novel Screening Method for Functional Disulfide-Rich Peptides by Solid-Phase Synthesis and Stabilization of mRNA–Protein Fusions. Nucleic Acids Research 2009, 37, e108, https://doi.org/10.1093/nar/gkp514.

20. Das, P.; Sercu, T.; Wadhawan, K.; Padhi, I.; Gehrmann, S.; Cipcigan, F.; Chenthamarakshan, V.; Strobelt, H.; dos Santos, C.; Chen, P.-Y.; et al. Accelerated Antimicrobial Discovery via Deep Generative Models and Molecular Dynamics Simulations. Nat. Biomed. Eng. 2021, 5, 613–623. https://doi.org/10.1038/s41551-021-00689-x.

21. Cardoso, M.H.; Orozco, R.Q.; Rezende, S.B.; Rodrigues, G.; Oshiro, K.G.N.; Cândido, E.S.; Franco, O.L. Computer-Aided Design of Antimicrobial Peptides: Are We Generating Effective Drug Candidates? Front. Microbiol. 2020, 10, https://doi.org/10.3389/fmicb.2019.03097.

22. Capecchi, A.; Zhang, A.; Reymond, J.-L. Populating Chemical Space with Peptides Using a Genetic Algorithm. J. Chem. Inf. Model. 2020, 60, 121–132. https://doi.org/10.1021/acs.jcim.9b01014.

23. Lu, H.; Zhou, Q.; He, J.; Jiang, Z.; Peng, C.; Tong, R.; Shi, J. Recent Advances in the Development of Protein–Protein Interactions Modulators: Mechanisms and Clinical Trials. Signal Transduct. Target Ther. 2020, 5, 213, https://doi.org/10.1038/s41392-020-00315-3.

24. Klebe, G. Virtual Ligand Screening: Strategies, Perspectives and Limitations. Drug Discov. Today 2006, 11, 580–594, https://doi.org/10.1016/j.drudis.2006.05.012.

25. Scior, T.; Bender, A.; Tresadern, G.; Medina-Franco, J.L.; Martínez-Mayorga, K.; Langer, T.; Cuanalo-Contreras, K.; Agrafiotis, D.K. Recognizing Pitfalls in Virtual Screening: A Critical Review. J. Chem. Inf. Model. 2012, 52, 867–881, https://doi.org/10.1021/ci200528d.

26. Amarasinghe, K.N.; De Maria, L.; Tyrchan, C.; Eriksson, L.A.; Sadowski, J.; Petrović, D. Virtual Screening Expands the Non-Natural Amino Acid Palette for Peptide Optimization. J. Chem. Inf. Model. 2022, 62, 2999–3007, https://doi.org/10.1021/acs.jcim.2c00193.

27. Friesner, R.A.; Banks, J.L.; Murphy, R.B.; Halgren, T.A.; Klicic, J.J.; Mainz, D.T.; Repasky, M.P.; Knoll, E.H.; Shelley, M.; Perry, J.K.; et al. Glide: A New Approach for Rapid, Accurate Docking and Scoring. 1. Method and Assessment of Docking Accuracy. J. Med. Chem. 2004, 47, 1739–1749, https://doi.org/10.1021/jm0306430.

28. Tubert-Brohman, I.; Sherman, W.; Repasky, M.; Beuming, T. Improved Docking of Polypeptides with Glide. J. Chem. Inf. Model. 2013, 53, 1689–1699, https://doi.org/10.1021/ci400128m.

29. Alogheli, H.; Olanders, G.; Schaal, W.; Brandt, P.; Karlén, A. Docking of Macrocycles: Comparing Rigid and Flexible Docking in Glide. J. Chem. Inf. Model. 2017, 57, 190–202, https://doi.org/10.1021/acs.jcim.6b00443.

30. Kurcinski, M.; Jamroz, M.; Blaszczyk, M.; Kolinski, A.; Kmiecik, S. CABS-Dock Web Server for the Flexible Docking of Peptides to Proteins without Prior Knowledge of the Binding Site. Nucl. Acids Res. 2015, 43, W419–W424, https://doi.org/10.1093/nar/gkv456.

31. Kurcinski, M.; Pawel Ciemny, M.; Oleniecki, T.; Kuriata, A.; Badaczewska-Dawid, A.E.; Kolinski, A.; Kmiecik, S. CABS-Dock Standalone: A Toolbox for Flexible Protein–Peptide Docking. Bioinformatics 2019, 35, 4170–4172, https://doi.org/10.1093/bioinformatics/btz185.

32. Porter, K.A.; Xia, B.; Beglov, D.; Bohnuud, T.; Alam, N.; Schueler-Furman, O.; Kozakov, D. ClusPro PeptiDock: Efficient Global Docking of Peptide Recognition Motifs Using FFT. Bioinformatics 2017, 33, 3299–3301, https://doi.org/10.1093/bioinformatics/btx216

33. Alam, N.; Goldstein, O.; Xia, B.; Porter, K.A.; Kozakov, D.; Schueler-Furman, O. High-Resolution Global Peptide-Protein Docking Using Fragments-Based PIPER-FlexPepDock. PLOS Comput. Biol. 2017, 13, e1005905, https://doi.org/10.1371/journal.pcbi.1005905.

34. Lamiable, A.; Thévenet, P.; Rey, J.; Vavrusa, M.; Derreumaux, P.; Tufféry, P. PEP-FOLD3: Faster de Novo Structure Prediction for Linear Peptides in Solution and in Complex. Nucl. Acids Res. 2016, 44, W449–W454, https://doi.org/10.1093/nar/gkw329.

35. Antunes, D.A.; Moll, M.; Devaurs, D.; Jackson, K.R.; Lizée, G.; Kavraki, L.E. DINC 2.0: A New Protein–Peptide Docking Webserver Using an Incremental Approach. Cancer Res. 2017, 77, e55–e57, https://doi.org/10.1158/0008-5472.CAN-17-0511.

36. Charitou, V.; van Keulen, S.C.; Bonvin, A.M.J.J. Cyclization and Docking Protocol for Cyclic Peptide–Protein Modeling Using HADDOCK2.4. J. Chem. Theory Comput. 2022, 18, 4027–4040, https://doi.org/10.1021/acs.jctc.2c00075.

37. Zhang, Y.; Sanner, M.F. AutoDock CrankPep: Combining Folding and Docking to Predict Protein–Peptide Complexes. Bioinformatics 2019, 35, 5121–5127. https://doi.org/10.1093/bioinformatics/btz459.

38. Tripathi, N.M.; Bandyopadhyay, A. High Throughput Virtual Screening (HTVS) of Peptide Library: Technological Advancement in Ligand Discovery. Eur. J. Med. Chem. 2022, 243, 114766, https://doi.org/10.1016/j.ejmech.2022.114766.

39. Jumper, J.; Evans, R.; Pritzel, A.; Green, T.; Figurnov, M.; Ronneberger, O.; Tunyasuvunakool, K.; Bates, R.; Žídek, A.; Potapenko, A.; et al. Highly Accurate Protein Structure Prediction with AlphaFold. Nature 2021, 596, 583–589. https://doi.org/10.1038/s41586-021-03819-2.

40. Baek, M.; DiMaio, F.; Anishchenko, I.; Dauparas, J.; Ovchinnikov, S.; Lee, G.R.; Wang, J.; Cong, Q.; Kinch, L.N.; Schaeffer, R.D.; et al. Accurate Prediction of Protein Structures and Interactions Using a Three-Track Neural Network. Science 2021, 373, 871–876. https://doi.org/10.1126/science.abj8754.

41. Baek, M.; Anishchenko, I.; Humphreys, I.R.; Cong, Q.; Baker, D.; DiMaio, F. Efficient and Accurate Prediction of Protein Structure Using RoseTTAFold2. bioRxiv 2023, https://doi.org/10.1101/2023.05.24.542179.

42. Evans, R.; O’Neill, M.; Pritzel, A.; Antropova, N.; Senior, A.; Green, T.; Žídek, A.; Bates, R.; Blackwell, S.; Yim, J.; et al. Protein Complex Prediction with AlphaFold-Multimer. bioRxiv 2022, https://doi.org/10.1101/2021.10.04.463034.

43. Humphreys, I.R.; Pei, J.; Baek, M.; Krishnakumar, A.; Anishchenko, I.; Ovchinnikov, S.; Zhang, J.; Ness, T.J.; Banjade, S.; Bagde, S.R.; et al. Computed Structures of Core Eukaryotic Protein Complexes. Science 2021, 374, eabm4805, https://doi.org/10.1126/science.abm4805.

44. Gulsevin, A.; Meiler, J. Benchmarking Peptide Structure Prediction with AlphaFold2. bioRxiv 2022. https://doi.org/10.1101/2022. 02.17.480937.

45. Tsaban, T.; Varga, J.K.; Avraham, O.; Ben-Aharon, Z.; Khramushin, A.; Schueler-Furman, O. Harnessing Protein Folding Neural Networks for Peptide–Protein Docking. Nat. Commun. 2022, 13, 176. https://doi.org/10.1038/s41467-021-27838-9.

46. Roney, J.P.; Ovchinnikov, S. State-of-the-Art Estimation of Protein Model Accuracy Using AlphaFold. Phys. Rev. Lett. 2022, 129, 238101, https://doi.org/10.1103/PhysRevLett.129.238101.

47. Anishchenko, I.; Pellock, S.J.; Chidyausiku, T.M.; Ramelot, T.A.; Ovchinnikov, S.; Hao, J.; Bafna, K.; Norn, C.; Kang, A.; Bera, A.K.; et al. De Novo Protein Design by Deep Network Hallucination. Nature 2021, 600, 547–552. https://doi.org/10.1038/s41586-021-04184-w.

48. Norn, C.; Wicky, B.I.M.; Juergens, D.; Liu, S.; Kim, D.; Tischer, D.; Koepnick, B.; Anishchenko, I.; Foldit Players; Baker, D.; et al. Protein sequence design using conformational landscape optimization. Proc. Natl. Acad. Sci. USA 2021, 118, e2017228118. https://doi.org/10.1073/pnas.2017228118.

49. Goverde, C.A.; Wolf, B.; Khakzad, H.; Rosset, S.; Correia, B.E. De Novo Protein Design by Inversion of the AlphaFold Structure Prediction Network. Protein Sci. 2023, 32, e4653, https://doi.org/10.1002/pro.4653.

50. Frank, C.; Khoshouei, A.; Stigter, Y. de; Schiewitz, D.; Feng, S.; Ovchinnikov, S.; Dietz, H. Efficient and Scalable de Novo Protein Design Using a Relaxed Sequence Space. bioRxiv 2023, https://doi.org/10.1101/2023.02.24.529906.

51. Kosugi, T.; Ohue, M. Solubility-Aware Protein Binding Peptide Design Using AlphaFold. Biomedicines 2022, 10, 1626, https://doi.org/10.3390/biomedicines10071626.

52. Rettie, S.A.; Campbell, K.V.; Bera, A.K.; Kang, A.; Kozlov, S.; De La Cruz, J.; Adebomi, V.; Zhou, G.; DiMaio, F.; Ovchinnikov, S.; Bhardwaj, G. Cyclic Peptide Structure Prediction and Design Using AlphaFold. bioRxiv 2023, https://doi.org/10.1101/2023.02.25.529

53. Mirdita, M.; Schütze, K.; Moriwaki, Y.; Heo, L.; Ovchinnikov, S.; Steinegger, M. ColabFold - Making Protein Folding Accessible to All. Nat. Methods 2022, 19, 679–682. https://doi.org/10.1038/s41592-022-01488-1.

54. Baek, M. (@minkbaek). Twitter Post: Adding a big enough number for “residue_index” feature is enough to model hetero-complex using AlphaFold (green&cyan: crystal structure / magenta: predicted model w/ residue_index modification). Available online: https://twitter.com/minkbaek/status/1417538291709071362 (accessed July 20, 2021).

55. Ovchinnikov, S. (@sokrypton). Twitter Post: Max Galettis alerted us to an error in our cyclic offset implementation. Which is now fixed in the notebook. (When you circularly permuted the sequences, the solutions were *nearly* identical (when aligned), but were not identical. Now with the bugfix, they are identical! With the bugfix, they are identical!) Available online: https://twitter.com/sokrypton/status/1670551262427840513 (accessed June 18, 2023).

56. Available online: https://github.com/sokrypton/ColabDesign/tree/main/af (accessed on 14 March 2022).

57. Zhang, Y.; Sanner, M.F. Docking Flexible Cyclic Peptides with AutoDock CrankPep. J. Chem. Theory Comput. 2019, 15, 5161–5168, https://doi.org/10.1021/acs.jctc.9b00557.

58. Word, J.M.; Lovell, S.C.; Richardson, J.S.; Richardson, D.C. Asparagine and Glutamine: Using Hydrogen Atom Contacts in the Choice of Side-Chain Amide Orientation. J. Mol. Biol. 1999, 285, 1735–1747. https://doi.org/10.1006/jmbi.1998.2401.

59. Morris, G.M.; Huey, R.; Lindstrom, W.; Sanner, M.F.; Belew, R.K.; Goodsell, D.S.; Olson, A.J. AutoDock4 and AutoDockTools4: Automated Docking with Selective Receptor Flexibility. J. Comput. Chem. 2009, 30, 2785–2791. https://doi.org/10.1002/jcc.21256.

60. Crooks, G.E.; Hon, G.; Chandonia, J.-M.; Brenner, S.E. WebLogo: A Sequence Logo Generator. Genome Res. 2004, 14, 1188–1190. https://doi.org/10.1101/gr.849004.

61. Stranges, P.B.; Kuhlman, B. A Comparison of Successful and Failed Protein Interface Designs Highlights the Challenges of Designing Buried Hydrogen Bonds. Protein Sci. 2013, 22, 74–82, https://doi.org/10.1002/pro.2187.

62. Adasme, M.F.; Linnemann, K.L.; Bolz, S.N.; Kaiser, F.; Salentin, S.; Haupt, V.J.; Schroeder, M. PLIP 2021: Expanding the Scope of the Protein-Ligand Interaction Profiler to DNA and RNA. Nucl. Acids Res. 2021, 49, W530–W534.

63. Banhos Danneskiold-Samøe, N.; Kavi, D.; Jude, K.M.; Nissen, S.B.; Wat, L.W.; Coassolo, L.; Zhao, M.; Asae Santana-Oikawa, G.; Broido, B.B.; Garcia, K.C.; Svensson, K.J. Rapid and Accurate Deorphanization of Ligand-Receptor Pairs Using AlphaFold. bioRxiv 2023, https://doi.org/10.1101/2023.03.16.531341.

64. Peccati, F.; Alunno-Rufini, S.; Jiménez-Osés, G. Accurate Prediction of Enzyme Thermostabilization with Rosetta Using AlphaFold Ensembles. J. Chem. Inf. Model. 2023, 63, 898–909, https://doi.org/10.1021/acs.jcim.2c01083.

65. Bryant, A.; Elofsson, A. EvoBind: in Silico Directed Evolution of Peptide Binders with AlphaFold. bioRxiv 2022, https://doi.org/10.1101/2022.07.23.501214.

66. Liu, J.; Chen, Z.; Li, Y.; Zhao, W.; Wu, J.; Zhang, Z. PD-1/PD-L1 Checkpoint Inhibitors in Tumor Immunotherapy. Front. Pharmacol. 2021, 12, https://doi.org/10.3389/fphar.2021.731798.

67. Yin, H.; Zhou, X.; Huang, Y.-H.; King, G.J.; Collins, B.M.; Gao, Y.; Craik, D.J.; Wang, C.K. Rational Design of Potent Peptide Inhibitors of the PD-1:PD-L1 Interaction for Cancer Immunotherapy. J. Am. Chem. Soc. 2021, 143, 18536–18547, https://doi.org/10.1021/jacs.1c08132.

