## Supplementary Materials for "Design of Cyclic Peptides Targeting Protein-Protein Interactions using AlphaFold"

### 1. SUPPLEMENTARY FIGURES

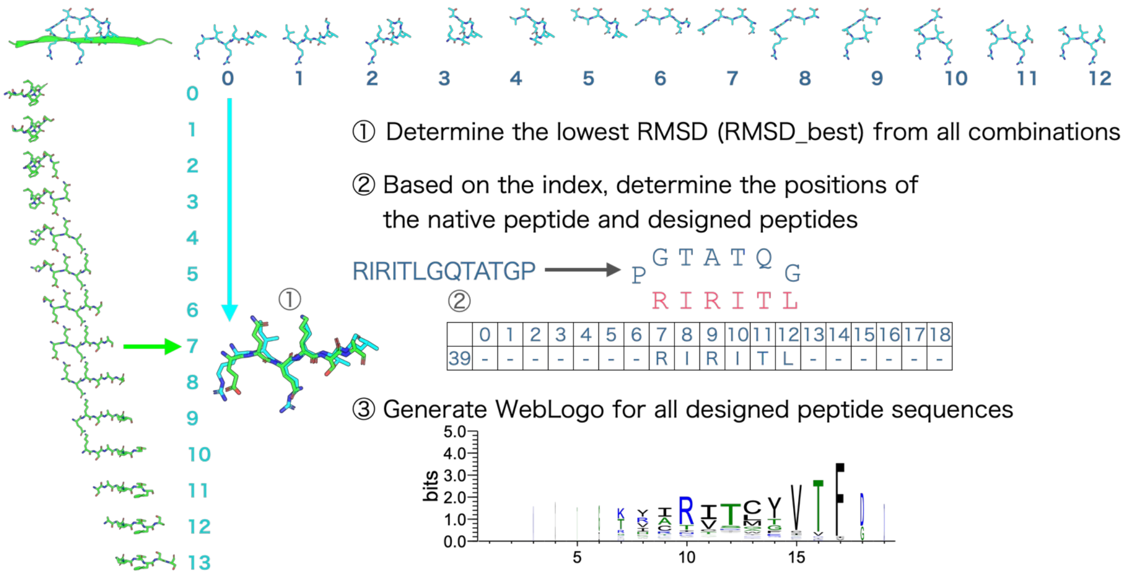


**Fig. S1.** Flowchart for the creation of sequence logos. At first, the RMSD was calculated for the cyclic peptide and native linear peptide C$\alpha$ of each window to determine the lowest RMSD of the C$\alpha$ among them. The lowest RMSD was named RMSD_best. Second, the design sequence was aligned based on the index numbers of the native peptide sequence and the cyclic peptide. After alignment, the non-6 letters were filled in with '-'. Third, sequence logos were created using WebLogo.

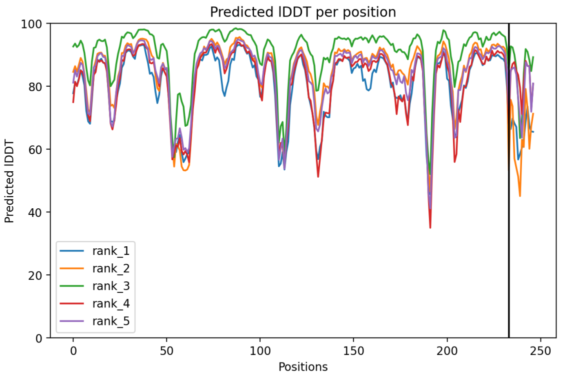


(**a**) AF2_ptm pLDDT

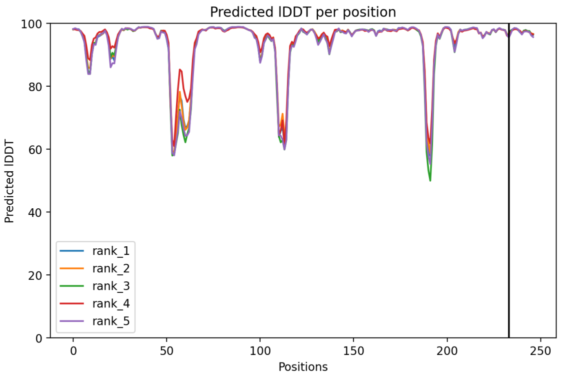


(**b**) AF2_v2 pLDDT

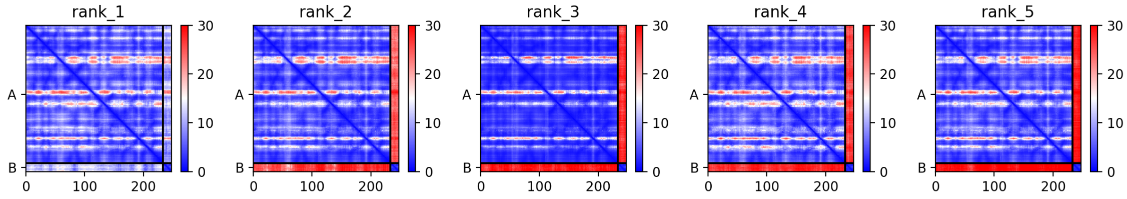


(**c**) AF2_ptm PAE

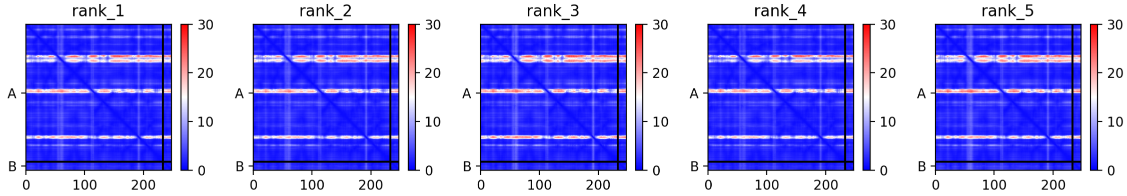


(**d**) AF2_v2 PAE

**Fig. S2.** Comparing the accuracy of the alphafold2_ptm(AF2_ptm) and alphafold2_multimer_v2 (AF2_v2) models for predicting protein-cyclic peptide complexes. **a**-**b** show the pLDDT results for 1SFI complex prediction using AF2_ptm and AF2_v2, respectively. **c**-**d** show the PAE results for 1SFI complex prediction using AF2_ptm and AF2_v2, respectively.

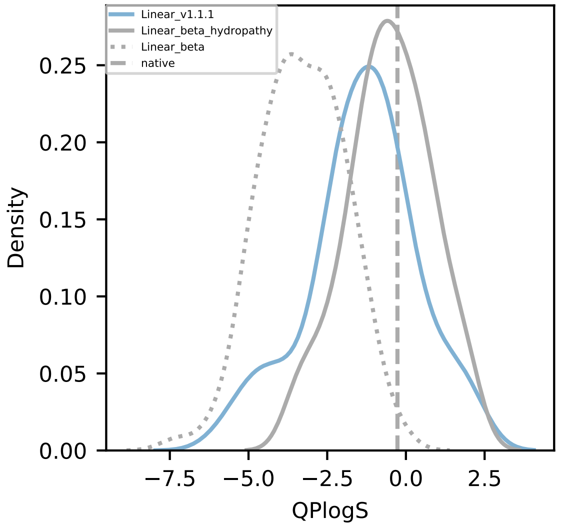


(**a**) QPlogS

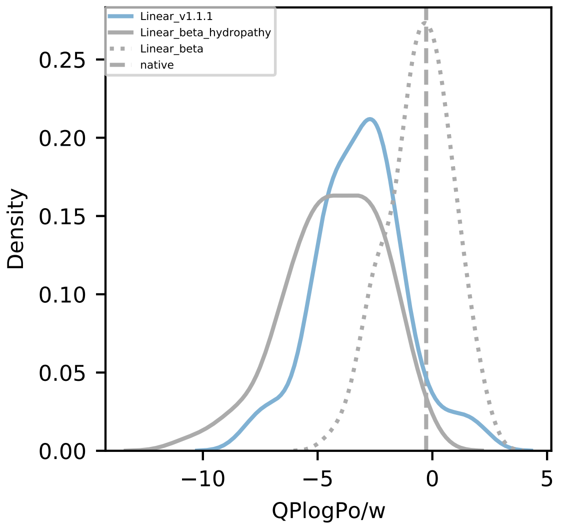


(**b**) QPlogPo/w

**Fig. S3.** Comparing the solubility and lipophilicity of the designed sequences in different versions of AfDesign. **a** and **b** show the QPlogS and QPlogPo/w results for the beta and v1.1.1 versions of AfDesign, respectively. The solid blue line shows the v1.1.1 result, the solid gray line shows the result of setting the weight of the hydropathy index to 0.5 in the beta version, the gray dotted line shows the result without the hydropathy index in the beta version (i.e., the default setting in the beta version), and the gray dashed line shows the result of the native p53 peptide.

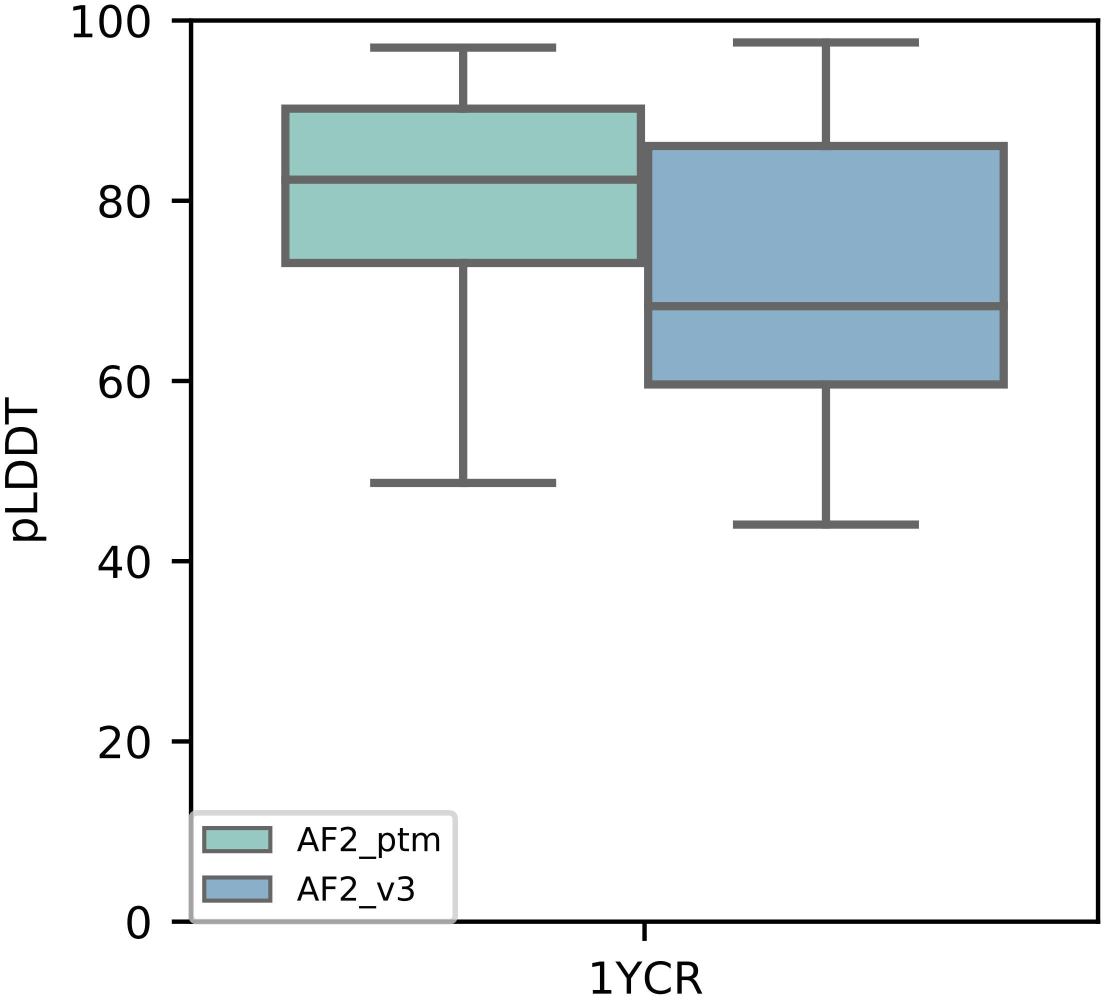


**Fig. S4.** Comparison of pLDDT for AF2_ptm- and AF_v3-designed sequences used in AfDesign. The pLDDT values for random sequences are the result of using AF2_ptm.

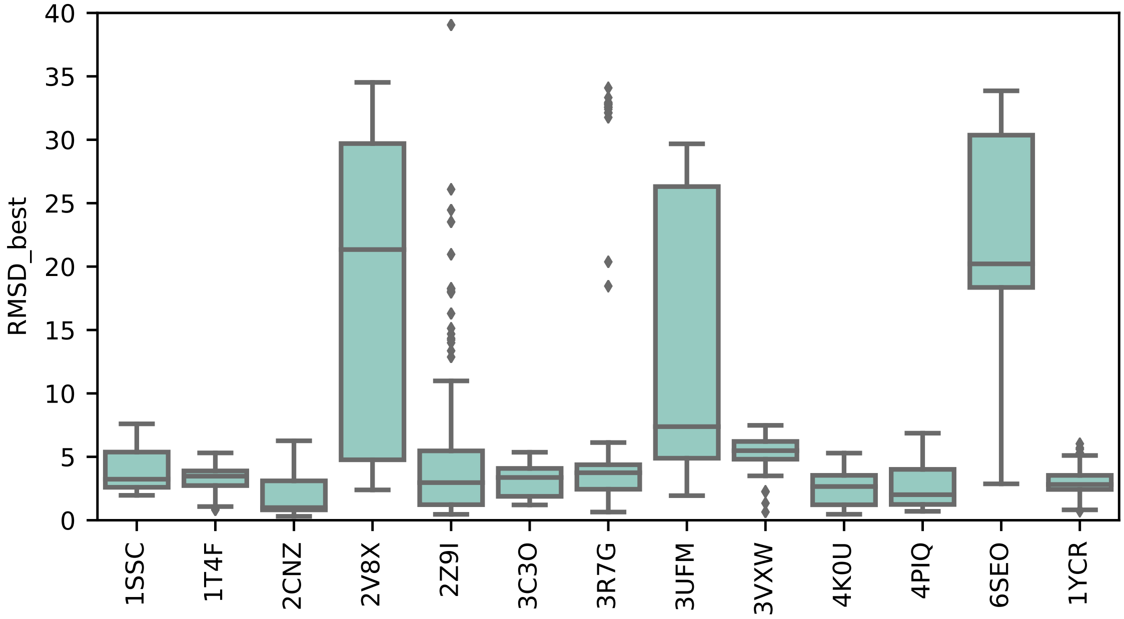


**Fig. S5.** Boxplots of RMSD_best for the predicted structures of the designed cyclic peptide and native peptides for 12 complexes and 1YCR.

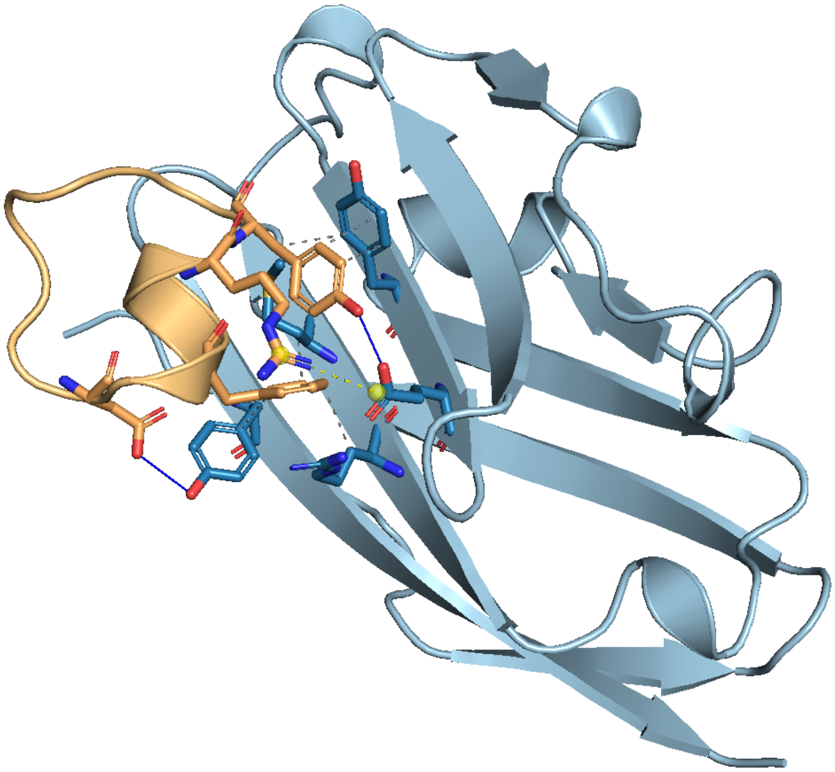

**Fig. S6.** Interatomic interaction analysis for PD-L1 with the designed cyclic peptide using PLIP. Interatomic interaction between PD-L1 and the designed peptide. The light blue color indicates PD-L1; light orange indicates the designed cyclic peptide. Hydrophobic bonds, hydrogen bonds, salt bridges, and charge centers are indicated using gray dashed lines, blue solid lines, yellow dashed lines, and yellow spheres, respectively.

### 2. SUPPLEMENTARY TABLES

**Table S1.** Correspondence between the PDB ID and chain of complexes applying cyclic binder hallucination in this study.

| PDB ID | protein chain | peptide chain |
|---|---|---|
| 1SSC | A | B |
| 1T4F | M | P |
| 1YCR | A | B |
| 2CNZ | A | B |
| 2V8X | A | B |
| 2Z9I | A | G |
| 3R7G | A | B |
| 3UFM | A | B |
| 3VXW | A | B |
| 4K0U | A | B |
| 4PIQ | A | B |
| 6SEO | A | L |
